# Titration of *in-cellula* affinities of protein-protein interactions

**DOI:** 10.1101/2020.04.27.063552

**Authors:** David Cluet, Blandine Vergier, Nicolas-Pierre Levy, Lucie Dehau, Alexandre Thurman, Ikram Amri, Martin Spichty

## Abstract

A genetic assay permits simultaneous quantification of two interacting proteins and their bound fraction at the single-cell level using flow cytometry. *In-cellula* affinities of protein-protein interactions can be extracted from the acquired data through a titration-like analysis. The applicability of this approach is demonstrated on a diverse set of interactions with proteins from different families and organisms and with *in-vitro* dissociation constants ranging from picomolar to micromolar.

---

The quest for methods that permit rapid and reliable determination of the affinity of protein-protein interactions (PPI) is unbroken. In contrast to biochemical *in-vitro* methods such as Isothermal Titration Calorimetry (ITC) and Surface Plasmon Resonance (SPR) that require purified proteins, quantitative genetic assays rely on the expression of the proteins of interest in cells. Many of these assays^1–6^ are inspired by the yeast two-hybrid (Y2H) technique^7–9^ which is based on the *in-cellula* expression of two proteins, usually named Bait and Prey, fused to an DNA-binding domain (BD) and an activation domain (AD), respectively. Upon physical interaction of the BD-Bait and AD-Prey proteins, a functional transcription factor is reconstituted that drives the expression of a reporter gene. The stronger the interaction, the higher should be the expression level of the reporter.^1^ However, the expression level of the AD-Bait and BD-Prey play an important role, too.^10^

We recently introduced a quantitative yeast-two hybrid system (qY2H) that permits for the first time simultaneous quantification of BD-Bait, AD-Prey and the reporter at the single-cell level without the need of any antibodies or purified proteins.^10^ Instead, we take advantage of fluorescent fusion proteins that can be detected by standard flow cytometers. Here we show how this qY2H method can be exploited to perform *in-cellula* affinity titrations by applying the following two important improvements:

1. Cellular contents of fluorescent proteins are determined in units of Molecules of Equivalent Soluble Fluorochrome (MESF), so that measured quantities become independent of the applied apparatus setup. It facilitates the future transferability of the qY2H measurements to other flow cytometers and allows researchers to consistently compare their results. Our reference fluorochrome is the yeast Enhanced Green Fluorescent protein (yEGFP) for which commercial calibration beads exist. The fluorescence intensity of TagRFP is converted into units of MESF of EGFP using independent calibration experiments with a fluorescent tandem protein BD-TagRFP-EGFP (see “Methods”).
2. We analyze the data by a titration-like procedure which allows the straightforward extraction of *in-cellula* dissociation constants for Bait:Prey interactions. In a proof of concept, we apply this *in-cellula* titration approach to a diverse set of PPIs with dissociation constants ranging from 117 pM to 17 μM (Table 1). As in *in-vitro* SPR experiments, each PPI can be measured by Y2H in two different orientations (by exchanging Bait and Prey). Here we study only the orientation that produced the higher reporter level.^10^ This orientation is considered as the molecular configuration with the higher accessibility of the PPI binding interface.^1^

**Table 1:**
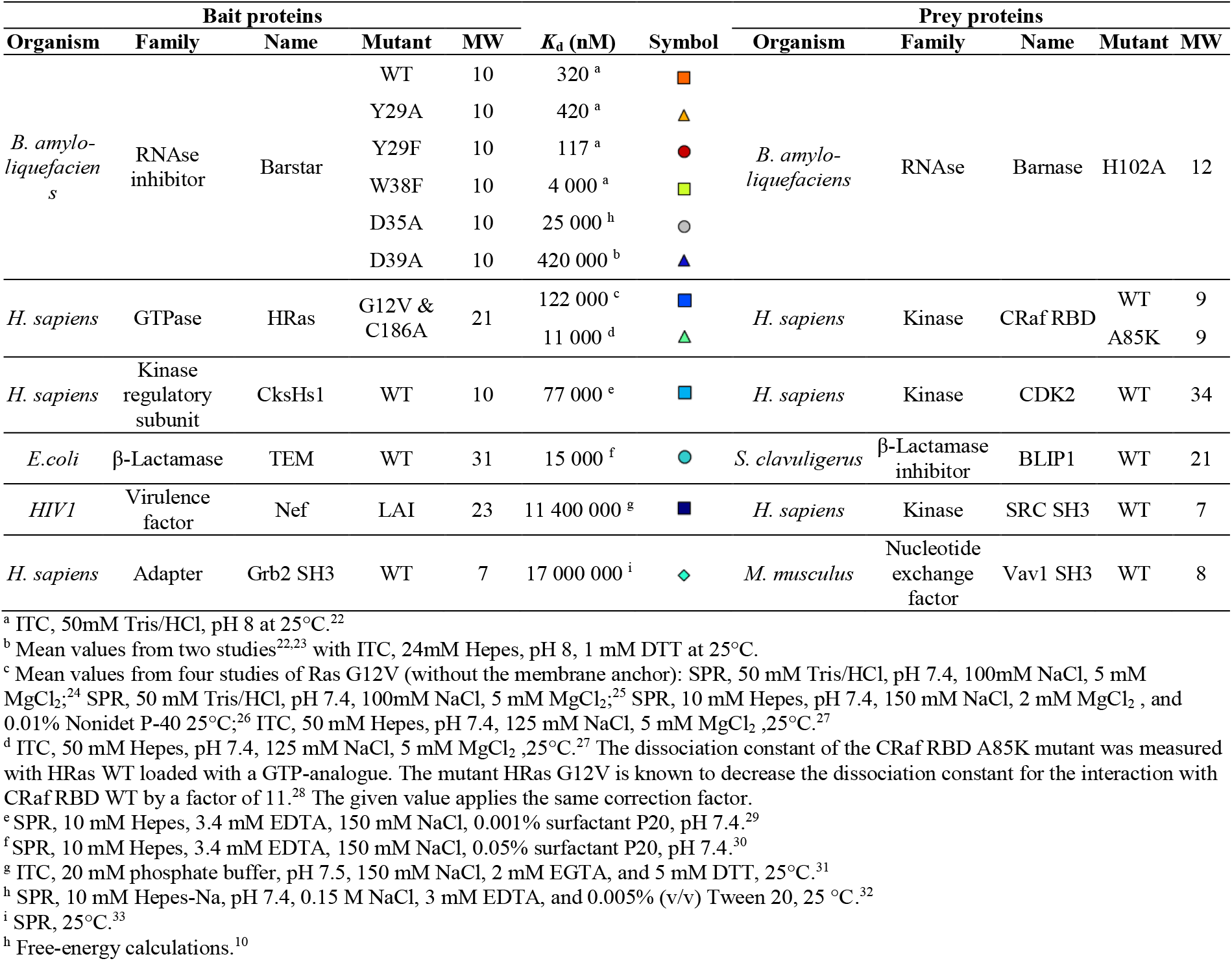
Investigated protein-protein interactions and their *in-vitro* affinities (*K*_d_).

In our qYH2 experiments, diploid yeast cells with constitutive expression of BD-Bait and induced expression of AD-Prey are cultured for two hours. Then, their fluorescence intensity is measured by flow cytometer in the three channels corresponding to TagRFP (BD-Bait), EGFP (AD-Prey), and TagBFP (reporter). Due to phenotypic variations, BD-Bait and AD-Prey are expressed at different levels among these cells which can be exploited to “prepare samples” for a titration. By gating, we can split the global heterogeneous ensemble of cells into several homogenous subensembles (bins). Each bin contains only cells within two specific, narrow intervals of red and green fluorescence intensity centered at values *R* and *G*, respectively. Assuming a linear relationship between fluorescence intensity and molecule numbers, *R* and *G* can be considered as measures for the mean cellular content of BD-Bait and AD-Prey in the corresponding bin.

With the mean value of the blue fluorescence intensity, we can calculate for each bin the normalized reporter level *φ*. It is obtained by forming the ratio of the expression level for the interaction of interest, *E*_interaction_ (Fig. 1A) and the level for a covalent BD-AD fusion, *E*_colvalent_ (Fig. 1B). This normalization renders *φ* dimensionless and independent of the acquisition apparatus (assuming again a linear relationship between molecule number and fluorescence intensity). Most importantly, we consider that *φ* reflects the time-averaged fraction of BD-Bait bound by AD-Prey during the reaction (as explained in the caption of Fig. 1). Thus, titration curves can be obtained when *φ* is plotted as a function of *G* while keeping *R* fixed (Fig. 1C). The curves can be fitted with the following Langmuir-type equation:

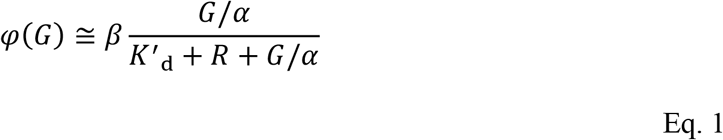

where *K*’_d_ is the *in-cellula* dissociation constant (in units of MESF of EGFP) and *α* and *β* are dimensionless parameters that empirically account for the fact that *φ* is a time-integrated property. The parameter *α* reduces the final cellular content of AD-Prey (measured at the end of the reaction, *G*) to the time-averaged content (over the entire reaction course, <*G*>). Since the induction kinetics under the GAL1-promotor in yeast^11^ displays a quadratic-like time dependence (for short induction times), a reasonable choice for *α* is 3 [<*G*> =_0_∫^1^ *Gt*^2^ *dt* = *G*/3]. The prefactor *β*, on the other hand, integrates differences in the expression kinetics of the reporter for *E*_interaction_ (induced expression) and *E*_covalent_ (constitutive expression). It can be determined experimentally by monitoring *φ* for *G* → ∞ using a high-affinity couple (such as BD-Barstar29F/AD-BarnaseH102A).

**Figure 1:**
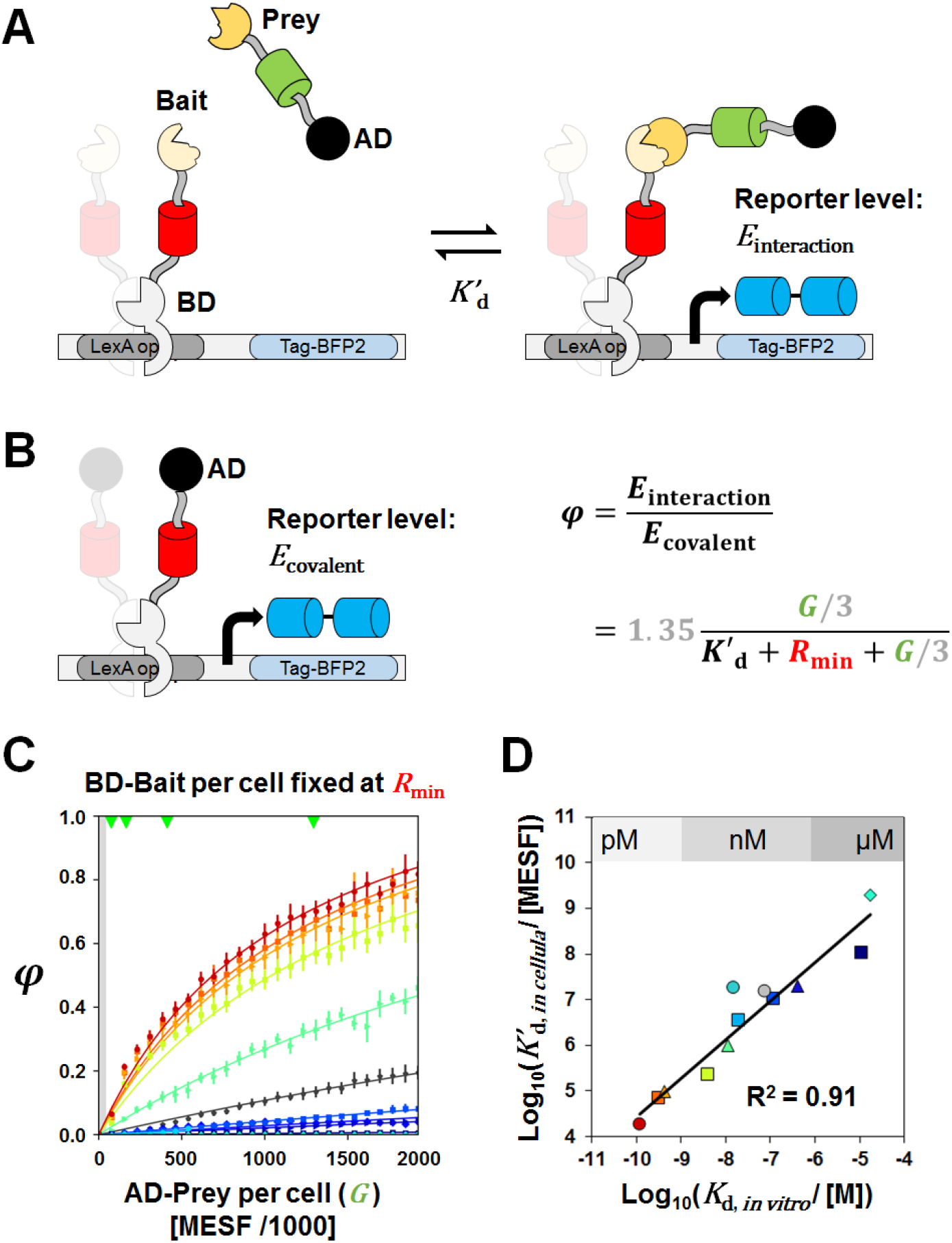
The qY2H system and its application to affinity titrations. (A) In our qY2H system, red-fluorescent BD-Bait interacts with green-fluorescent AD-Prey to reconstitute a transcription factor that drives the expression of a blue-fluorescent reporter. Our hypothesis is that the expression level of the reporter, *E*_interacting_, reflects the number of BD-molecules bound to the promotor corrected by the fraction of BD-Bait bound to AD-Prey. This fraction is influenced by the affinity between the Bait and the Prey, but also by the expression levels of BD-Bait and AD-Prey. (B) When the activation domain is covalently linked to the DNA-binding domain, the expression level *E*_covalent_ depends only on the number of BD-Bait molecules bound the promotor. Thus, when forming the quantity *φ* by dividing *E*_interacting_ with *E*_covalent_, we obtain a measure for the fraction of BD-molecules bound by an AD-Prey molecule. To determine *E*_covalent_, we constructed a BD-AD fusion protein. Unfortunately, the activation domain B42 (as used in **a**) turned out to be toxic for our yeast strains when used in the BD-AD construct. Instead, we used the activation domain B112. Difference in the activation potential between B42 and B112 are integrated in the parameter *β* of Eq. 1. (C) The quantity φ can be monitored as a function of different levels of EGFP Molecules of Equivalent Soluble Fluorochrome (MESF) corresponding to different cellular levels of AD-Prey. In these titrations, the level of BD-Bait is kept fixed at the lowest possible value (see “Methods”). For the interaction TEM/BLIP1 (cyan line) the titration can be preformed only up to one third of the titrant quantity due to expression problems of AD-BLIP1.^10^ Green triangles at the top vertical axis indicate the position of used calibration beads. (D) When the titration curves are fitted with Eq. 1, we can extract the dissociation constant in units of MESF (*K*’_d_). The estimated *K*’_d_-values show a remarkable correlation with the dissociation constants measured from alternative *in-vitro* experiments (with matching symbols of Table 1).

We recommend that the titrations are carried out with the lowest possible value of *R*=*R*_min_ (as defined by the detection limit of TagRFP by flow cytometery, see “Methods”). It limits overexpression and associated protein burden effects.^12^ Furthermore, the auto-activation potential of the BD-Bait fusion is kept at a minimum, too.^13^ Most importantly, it mimics the condition of *in-vitro* affinity titration experiments^14^ where the concentration of the titrated species (here BD-Bait) is kept fixed and as low as possible to avoid saturation effects. For the titrations with *R*=*R*_min_ the parameters *α*=3 and *β* =1.35were used to extract the *K*’_d_-values.

Despite substantial differences between our *in-cellula* system and *in-vitro* setups (as previously discussed^10^ in detail), the *in-cellula* affinities strongly correlate with those from *in-vitro* measurements (R^2^=0.91, Fig. 1D). The slope of the regression line is 0.84. Other *in-cellula* assays usually find lower correlation coefficients (< 0.9) and significantly lower values for the slope of the regression line (0.2-0.6).^4,5,15–17^ This is even more remarkable if one considers that the tested set of PPIs in this work is significantly more diverse. It may indicate a higher sensitivity for the qY2H titration approach; more testing will be necessary to confirm this surmise.

The presented protocol is robust as witnessed by the small error bars in the titration curves (Fig. 1c). All steps of the protocol have been optimized in liquid phase that can be easily automated for the use of microplates and integrated within robotic pipelines. It sets the stage for high-throughput affinity screenings of PPIs using cross-mating approaches^18,19^ with libraries of yeast clones. As an outlook, affinity-based networks^20^ can be created by attributing weights to the PPI edges according to their effective affinities. It contrasts standard Y2H screens that yield networks with only binary information (YES or NO). The topology of edge-weighted spring-embedded networks^21^ may help identifying key pathways within the network, and how they change as a function of environmental conditions (stress, metabolism, *etc*). Thus, we anticipate that high-throughput qY2H affinity data would boost the modelling of interactomes and thereby advance significantly systems biology.

## Methods

The qY2H experiments, acquisitions by flow cytometry and analyses were carried out as described in our previous study10 with the following particularities. About 107 cells were cultured per experiment and interaction (including the covalent BD-AD fusion and the control sample BD-Empty / AD-Empty). To ensure that these cells have been indeed transfected with all three vectors, we selected for the analysis large (=growing) cells with a forward scatter range 75 000 < FSC-H < 125 000; “H” indicates signal height. Furthermore, only cells with a red fluorescence intensity of 800 ± 100 TagRFP-H were analysed. This bin is located just above the 95% threshold of the non-fluorescent cells,^10^ and therefore defines *R*_min_.

The mean Tag BFP-H value was then calculated for bins of varying *G* values from −500 to 25500 yEGFP-H (bin size 1000). For each bin we calculated:

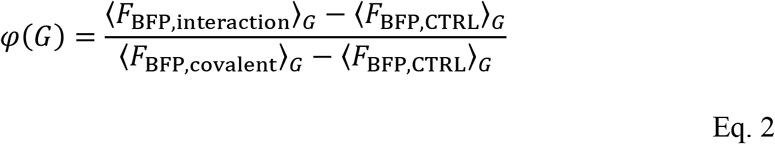

where <*F*_BFP,X_>_*G*_ is the mean blue fluorescence intensity. The subscripted X refers to the physical interaction, covalent fusion or control couple. The control couple BD-Empty / AD-Empty^10^ permits to remove the background of the reporter system.

Finally, *G* values were converted into MESF of EGFP using calibration beads (Ozyme, reference 632594) following the manufactor’s protocol. For the conversion of *R*_min_, we performed independent calibration measurements with diploid yeast cells expressing the fluorescent tandem fusion protein BD-TagRFP-yEGFP (under the same condition as the qY2H experiments). Cells with a red fluorescence intensity of *R*_min_ = 800 ± 100 TagRFP-H displayed a mean green fluorescence intensity of 370 000 MESF of EGFP.

Experiments and analyses were performed at least three times for each interaction and averaged titration curves were least-square fitted with Eq. 1.

## Acknowledgments

This work was supported by the Fonds Recherche of the Ecole Normale Supéreure de Lyon.

